# sincFold: end-to-end learning of short- and long-range interactions in RNA secondary structure

**DOI:** 10.1101/2023.10.10.561771

**Authors:** Leandro A. Bugnon, Leandro Di Persia, Matias Gerard, Jonathan Raad, Santiago Prochetto, Emilio Fenoy, Uciel Chorostecki, Federico Ariel, Georgina Stegmayer, Diego H. Milone

## Abstract

**Motivation:** Coding and non-coding RNA molecules participate in many important biological processes. Non-coding RNAs fold into well-defined secondary structures to exert their functions. However, the computational prediction of the secondary structure from a raw RNA sequence is a long-standing unsolved problem, which after decades of almost unchanged performance has now re-emerged thanks to deep learning. Traditional RNA secondary structure prediction algorithms have been mostly based on thermodynamic models and dynamic programming for free energy minimization. More recently deep learning methods have shown competitive performance compared with the classical ones, but still leaving a wide margin for improvement.

**Results:** In this work we present sincFold an end-to-end deep learning approach that predicts the nucleotides contact matrix using only the RNA sequence as input. The model is based on 1D and 2D residual neural networks that can learn short- and long-range interaction patterns. We show that structures can be accurately predicted with minimal physical assumptions. Extensive experiments were conducted on several benchmark datasets, considering sequence homology and cross-family validation. sincFold was compared against classical methods and recent deep learning models, showing that it can outperform state-of-the-art methods.

**Availability:** The source code is available at https://github.com/sinc-lab/sincFold (v0.16) and the web access is provided at https://sinc.unl.edu.ar/web-demo/sincFold

## 1. Introduction

Non-coding ribonucleic acid (ncRNA) molecules have emerged as crucial players in cellular processes, encompassing epigenetics, transcriptional and post-transcriptional regulation, chromosome replication, translation and protein activity and stability [31, 55]. Recent efforts have even explored the clinical potential of ncRNA in diagnostics, vaccines, and therapies [51]. This paradigm shift in our understanding of ncRNA, from being dismissed as “transcriptional noise” prior to the 1980s to being recognized as regulators of gene expression at multiple levels, has generated an explosion of research in this field over the past few decades [9].

RNA itself consists of an ordered sequence of four basic nucleotides: adenine (A), cytosine (C), guanine (G), and uracil (U). Pairing these bases within an RNA molecule gives rise to its secondary structure, a crucial determinant of its functions and stability [43]. The secondary structure is characterized by hydrogen bonding interactions between complementary base pairs, which typically include the canonical Watson-Crick-Franklin pairs A-U and C-G [50], along with the wobble pair G-U [48]. Basic stem-loop structures, formed by nested base pairs, are commonly observed. However, the secondary structure can also exhibit complex motifs arising from local bonding and long-range sequence interactions.

Despite the growing number of publicly accessible ncRNA sequences, a significant proportion of their true structures remains unknown [17]. Secondary structures can be obtained with sophisticated experimental techniques such as X-ray crystallography, nuclear magnetic resonance [16, 24, 44], enzymatic probing methods such as nextPARS [10], or chemical probing such as DMS-seq [11] and SHAPE-seq [26]. However, all these methods suffer from low resolution and high costs [36]. Consequently, due to its cost-effectiveness the computational prediction has gained substantial relevance in biological research and biotechnological applications.

Traditional computational methods for RNA secondary structure prediction employ a thermodynamic model of base-pair interactions optimized through dynamic programming to identify structures with minimal free energy [40, 47, 57]. Despite being proposed 20 years ago [30], these methods, such as RNAstructure [34], ProbKnot [5], RNAfold [25], LinearFold [21], LinearPartition [54], and Ipknot [38], continue to dominate the field. However, their average base-pair prediction performance remains at around *F*_1_ = 70% [7].

To surpass this performance ceiling, machine learning (ML) techniques, and particularly deep learning (DL), have emerged as promising alternatives [52]. DL techniques have been widely noticed for structure prediction in proteins with AlphaFold [22] and more recently several methods were presented for RNA secondary structure prediction [8, 15, 38, 41, 53]. However the available RNA datasets are very small compared to proteins, they are highly biased in several ways and pseudoknots are not consistently annotated, being a key factor in RNA structures. In [39] authors state that there are several possible ways to enable the accurate prediction of RNA structures in the near future, such as improving knowledge through more data, diversifying the data used in prediction, and improving the machine learning methods used. In particular DL methods rely less on assumptions about the thermodynamic mechanics of folding, instead adopting a data-driven approach. Consequently, they could be better suited to identify complex structures that defy modeling using traditional techniques. However, recent systematic evaluations of techniques for comparatively assessing their performance on ncRNAs showed that DL has not yet clearly overperformed classical methods [7, 12, 23].

Currently, several DL approaches are available with different architectural designs, input representations, training data and optimization algorithms for parameter adjustments [56]. Among these proposals, SPOT-RNA [41] was the pioneering DL method based on ensembles of convolutional networks (CNNs) and bidirectional Long-short term memory neural networks (LSTM). SPOT-RNA2 [42] improved its predecessor by using predictions from thermodynamic models, evolution-derived sequence profiles and mutational coupling, however requiring multiple sequence alignments. Another hybrid approach was MXfold [1], combining support vector machines and thermodynamic models. Similarly, DMFold [49] and MXFold2 [38] integrated DL techniques with energy based methods. Another method based on both DL and dynamic programming was CDPfold [53], which iteratively computes a matrix representation of possible matchings between bases according to a physical model of base interactions, and then trains a convolutional network over this matrix to predict base pairing probabilities. Upon this, dynamic programming is applied to obtain the final RNA secondary structure.

In more recent years, UFold [15] approached the secondary structure prediction problem using a well-known architecture from image segmentation, the U-Net encoder-decoder [35]. It uses a 2D feature map to encode the occurrences of one of the sixteen possible base pairs between nucleotides for each position in the map, including an additional channel with the matrix representation of possible matchings iteratively computed with the algorithm proposed in CDPfold. The predicted output is the contact score map between the bases of the input sequence, which goes through a post-processing step that involves solving a linear programming problem to obtain the optimum contact map. Interestingly, a very recent method, REDfold [8], reported to outperform UFold. This DL method also utilizes a U-Net encoder-decoder network to learn dependencies among the RNA sequence, together with symmetric skip connections to propagate activation information across layers and output post-processing with constrained optimization.

In this work we present sincFold, a novel end-to-end deep learning method for RNA secondary structure prediction. Our approach is based on ResNet bottlenecks to capture both short- and long-range dependencies in the RNA sequence. Unlike other DL models, we adopt a two-stage encoding process: initially, we model sequence encoding in 1D, enabling the learning of small context features and reducing computational costs; then a pairwise encoding in 2D is incorporated to capture distant relationships. Extensive experimental evaluations on two widely used ncRNA databases demonstrate that sincFold outperforms classical methods and DL state-of-the-art techniques in terms of *F*_1_ performance. We have made the source code for sincFold freely accessible, facilitating its adoption and further development in the research community^1^. Moreover, a web service to test the trained model is provided^2^.

## 2. The sincFold model

In order to obtain a secondary structure prediction from a stand-alone RNA sequence, we propose sincFold. As shown in Figure 1, this novel deep learning architecture is composed of two stages: the first one learns local patterns in 1D encodings, while the second stage can learn more distant interactions in 2D. The figure represents in detail the shapes and dimensions of the data along the pipeline (top) and the neural processing blocks (bottom).

**Figure 1:**
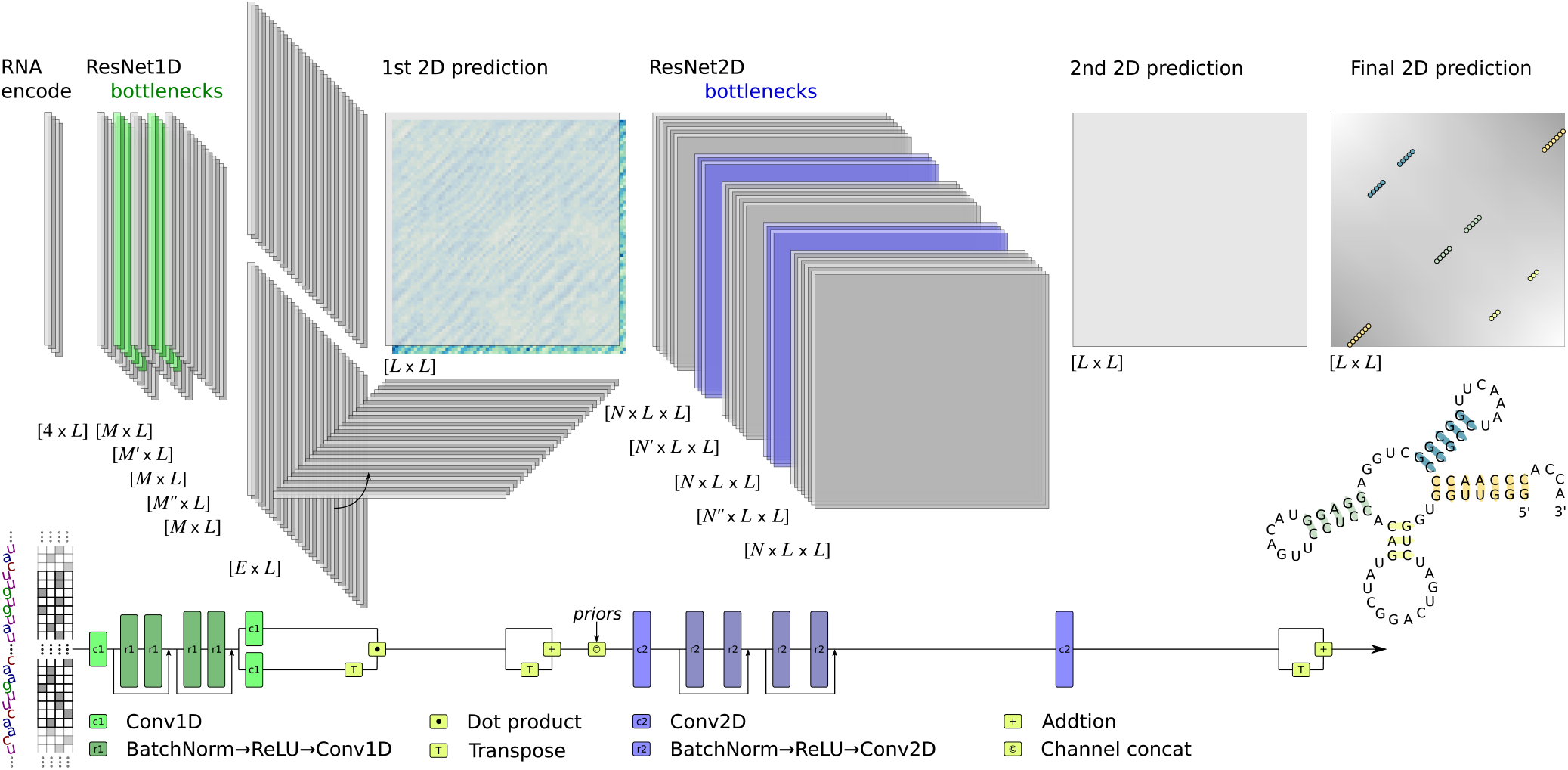
The end-to-end architecture of sincFold. Top: data flow with its shapes and dimensionality in each point of the architecture, from the [4*×L*] one-hot encoded RNA sequence at the input to the [*L×L*] connection matrix at the output. Bottom: neural processing blocks depicted as differentiable layers.

The model takes as input a RNA sequence of length L encoded in one-hot (bottom-left) so that each nucleotide type of the sequence is represented with a vector of size 4 (i.e. a one-hot codification of the 4 canonical nucleotides). The encoded sequence goes through a one-dimensional (1D) convolutional layer that performs a first automatic extraction of low-level features for each nucleotide. Then, identity blocks [19] are stacked in a 1D-ResNet. These blocks allow the model to propagate the signal and reduce vanishing gradient issues, while maintaining the same sequence length. Moreover, the identity blocks make the model capable of auto-defining the number of convolutional layers needed during training. Each block is composed of two batch normalization layers, ReLU activations and convolutional layers in 1D, with bottlenecks in the features (depicted with light green in the figure). Bottlenecks reduce the learnable parameters while helping to learn more relevant features.

After the 1D bottlenecks, a *M × L* encoding is obtained, where *M* is the dimension of the feature vector of each nucleotide. Then two convolutions in 1D produce two compressed encodings of size *E × L*. A matrix product between one *E × L* matrix and the other *E × L* matrix transpose is made, obtaining a first “draft” of the contact matrix in 2D (*L × L*). After that, the matrix is forced to be symmetric by adding its transpose. An additional channel of interaction priors is added at this point, coding different bonding strengths for C-G, U-A and G-U.

Once the information is represented in 2D, the new tensor *L×L* will go through a 2D-ResNet stage. Similarly to the 1D-ResNet stage, a 2D-convolutional layer is followed by 2D-ResNet blocks composed of batch normalization layers, ReLU activations and 2D convolutions. After several 2D-ResNet layers with bottlenecks the 2D pairwise encodings are flattened to a *L × L* output, and its transpose is added to force symmetry. This output matrix is the final 2D prediction of the secondary structure for the RNA sequence, and the entire model can be trained with a unified cost function. A simple post-processing is applied to find the maximum activation on each row and column, thus retaining only one interaction per nucleotide.

To guide training, we propose a composed loss function

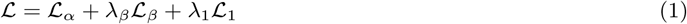

where *ℒ*_*α*_ is the cross-entropy loss of the final prediction, *ℒ*_*β*_ is the cross-entropy loss of the model prediction prior to the 2D-ResNet block, and *ℒ*_1_ is a *L*_1_ loss of the predictions used to enforce the contact matrices sparseness. Cross-entropy is computed element by element in the matrix. The weights *λ*_*β*_ and *λ*_1_ of each of these terms are hyperparameters to be adjusted experimentally.

Our proposed architecture is different to existing DL models in several ways. SPOT-RNA converts to a 2D representation but only as a pre-processing stage, by outer concatenation of the one-hot codification of the sequence. In this model, 1D patterns are not learned throughout the sequence. Then, the prediction of structure is obtained with an ensemble of ResNet blocks with dilated convolutions, a 2D-BLSTM (bidirectional long short-term memory) layer and a fully connected block. Furthermore, the SPOT-RNA source code for training is not available and thus it cannot be compared to other methods under the same conditions. MXFold2 has an architecture that models 1D and 2D representations though BLSTM and 2D convolution blocks, respectively. The conversion from 1D to 2D is based on a concatenation of halves of the 1D embeddings, so that different halves appear together in the corresponding coordinates of the *L × L* output. The choice of halves as a concatenation block only responds to the need to form a 2D representation, but has no basis in the modeling of structural connections to be predicted. Moreover, as in SPOT-RNA pre-processing, these concatenations do not include any inner products that measure similarity between 1D representations. Finally, it is important to note that MXFold2 is actually a hybrid method, which does not predict a contact matrix but four types of folding scores for each pair of nucleotides. The folding scores are integrated with the free energy parameters of Turner nearest-neighbor model. Then an optimal secondary structure is calculated using classical dynamic programming. Differently from UFold and REDfold, which use the standard U-Net originally proposed in computer vision for image segmentation, and a post-processing step with linear programming, with sincFold we propose a novel full end-to-end architecture that models separately the 1D (short range) and 2D (long range) interaction. It is important to note that in both UFold and REDfold the conversion from 1D to 2D is, as in other models, a pre-processing stage (i.e. it is not part of the DL model). For example, in UFold pre-processing the one-hot codification of the RNA sequence is converted into a 16 channels ‘image’ via a Kronecker product. In contrast, sincFold learns representations from a 1D sequence, converts them to a 2D representation with a tensorial product and then learns long range interactions through training.

## 3 Data and performance measures

### 3.1 Data

In order to evaluate the performance of sincFold, we have run benchmarks with datasets widely used by the community.

#### RNAstralign dataset [46]

contains 37,149 sequences from 8 large RNA families: 5S rRNAs, Group I Intron, tmRNA, tRNA, 16S rRNA, Signal Recognition Particle (SRP) RNA, RNase P RNA and Telomerase RNA. It is one of the most comprehensive RNA structure datasets available.

#### ArchiveII dataset [43]

the most widely used benchmark dataset for RNA folding methods, containing RNA structures from 9 RNA families: 5S rRNAs, SRP RNA, tRNA, tmRNA, RNase P RNA, Group I Intron, 16S rRNA, Telomerase RNA and 23S rRNA. The total number of sequences is 3,975.

#### TR0-TS0 dataset [41]

the same train and test sets as in SPOT-RNA. The learning data are from bpRNA 1.0 (Danaee et al., 2018). It consists in a nonredundant set of RNA sequences with annotated secondary structure from bpRNA34 at 80% sequence-identity cutoff with CD-HIT-EST [14]. This filtered dataset of 13,419 RNAs is randomly divided into 10,814 RNAs for training (TR0), 1,300 for validation (VL0), and 1,305 for an independent test (TS0).

#### Ablation dataset

in addition, we compiled a dataset of sequences derived from the URS server [4] to be used as a small independent dataset for model optimization. These sequences and secondary structures were extracted from the Protein Data Bank (PDB), consisting of 753 sequences ranging from 8 to 456 nucleotides.

As suggested in [45], sequences longer than 512 nucleotides were filtered to limit the runtime of experiments, leaving 22,611 sequences in the RNAstralign dataset and 3,864 sequences in the ArchiveII dataset. Group I intron RNAs were excluded from the RNAstralign dataset because it included sequences without a unique structure. Thus, in this manuscript we will show results only for sequences with less than 512 nt.

To assess the performance, all DL methods used in this study were re-trained from scratch with the exact same partitions for training and testing. First we perform a *k*-fold cross-validation with *k* = 5 on the Ablation, ArchiveII and RNAstralign datasets. For the ArchiveII dataset, the original *k*-fold split provided by the authors was used [43]. Sequences were randomly divided into 5 independent folds of approximately the same size, and each fold was in turn taken as the test data while the remaining folds were taken as the training data. Then, we considered the structural differences between sequences used in training and testing, in order to analyze the impact of homology on performance. In the TR0-TS0 dataset, we use the provided homology-aware partitions with 80% sequence similarity cut-off. Finally, we perform a cross-family analysis (testing on unseen RNA families) using the ArchiveII dataset.

### 3.2 Performance measures

The focus of performance measures is on the predicted base pairs in comparison to a reference structure [29]. Pairs that are both in the prediction and in the reference structure are true positives (TP), while pairs predicted but not in the true structure are false positives (FP). Similarly, a pair in the reference structure that is not predicted is a false negative (FN), and a pair that is neither predicted nor in the true structure is a true negative (TN). To fully characterize the successes and failures of structure prediction, we use the *F*_1_ score, defined as

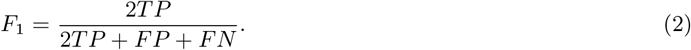

The whole RNA structure can be considered as a large interaction network composed of interactions and base stackings [27]. The interaction network fidelity (INF) similarity measure [32] was designed to score the similarity between the interactions of a reference RNA structure and the interactions of a predicted RNA structure. INF is defined as

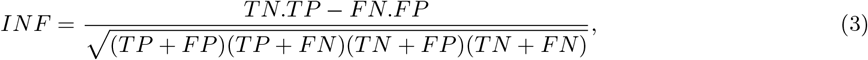

resembling the well-known Matthews correlation coefficient. When the prediction reproduces exactly the base interactions of the reference structure, then |*FP* | = |*FN* | = 0, |*T P* | *>* 0, and thus *INF* = 1. When the prediction does not reproduce any of the interactions of the reference structure, then *INF* = 0, since |*T P* | = 0.

In [37] it was demonstrated how two structures that share a common feature (for example, a hairpin) with the exact same base pair patterns can achieve *F*_1_ = 0. This is because similar base-pair patterns between the two secondary structures can only be shifted, but this will not be reflected by the *F*_1_ score. Thus, the Weisfeiler-Lehman graph kernel (WL) metric was proposed in order to capture graphs structural information by iteratively refining node labels based on their local neighborhoods. The WL metric first assigns to each node (nucleotide) in the graph (secondary structure) a label representing its local structural information. Then, a label propagation step iterates over the nodes and updates their labels based on the labels of their neighboring nodes. Finally, a hash function is computed that aggregates these labels to generate a feature vector. The WL is defined as

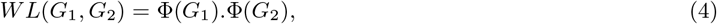

where Φ(*G*_*i*_) represents the feature vector of graph *G*_*i*_ obtained by aggregating the labels through the hash functions. The WL-similarity score is sensitive to both structural and sequence-level alterations.

### 3.3 Distance measure for secondary structures

It is known that minor changes in RNA sequences can represent significant changes in secondary structure and, conversely, very similar structures can be obtained from quite different sequences. Thus, for analyzing results, the structural distance between data samples is more representative of the prediction challenge than a simple sequence-level distance. For this reason, the structural distance was computed using RNAdistance from the ViennaRNA package [13, 25]. This distance is based on the edit distance of a tree representation, in which the secondary structure is converted into a tree by assigning an internal node to each base pair and a leaf node to each unpaired digit [20]. Then, a tree is transformed into another tree by a series of editing operations with predefined costs. The distance between the two trees is the smallest sum of the costs along an editing path, which is divided by the length of the longest sequence in order to obtain a normalized distance.

## 4 Results

### 4.1 Ablation study and hyperparameters exploration

We conducted an ablation study to gain a deeper understanding on the contribution of each of the components of the sincFold architecture. We run several versions of the sincFold model: C1D) a baseline model with only 1D-convolutional networks; R1D) the same model replacing convolutions with 1D-residual blocks and bottlenecks; C1D+C2D) the model with the 2D-stage using only convolutional neural networks; C1D+R2D) replacing convolutions with the 2D-residual blocks in the 2D stage; and R1D+R2D) with residual blocks and bottlenecks in both stages. The *F*_1_ scores for each ablated sincFold version, from a 5-fold cross-validation on the Ablation dataset, are shown in the boxplots of Figure 2.

**Figure 2:**
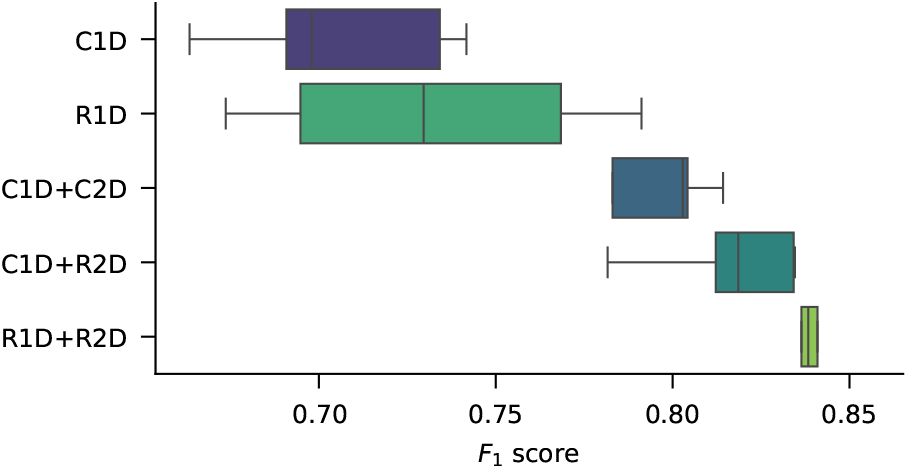
Ablation study on each of the components of the sincFold architecture. Each box has the *F*_1_ scores from a 5-fold cross-validation on the Ablation dataset.

It can be seen that changing the C1D to a R1D block slightly improves the median results, from a median *F*_1_ = 0.697 to *F*_1_ = 0.729. It can be observed that adding the 2D stage (C2D) to the output of the previous models increased their performance significantly, by 10% for each model. The *F*_1_ raises up to 0.802 with C1D+C2D, and using a ResNet instead of a CNN in the 2D stage (C1D+R2D), performance further improves up to *F*_1_ = 0.818. Finally, results are even further improved in the model with ResNet blocks in both stages (R1D+R2D), reaching *F*_1_ = 0.838.

After the ablation study we conclude that ResNet blocks effectively improve the generalization capability, in comparison to simple convolutional layers. Figure 3 presents a detailed analysis of the true positive rate of predictions in this dataset along the interconnection distance. Interestingly, when the 2D stage (green) is added to the 1D stage (blue) the model performance improves for all connections, and especially for distances longer than 200 nt.

**Figure 3:**
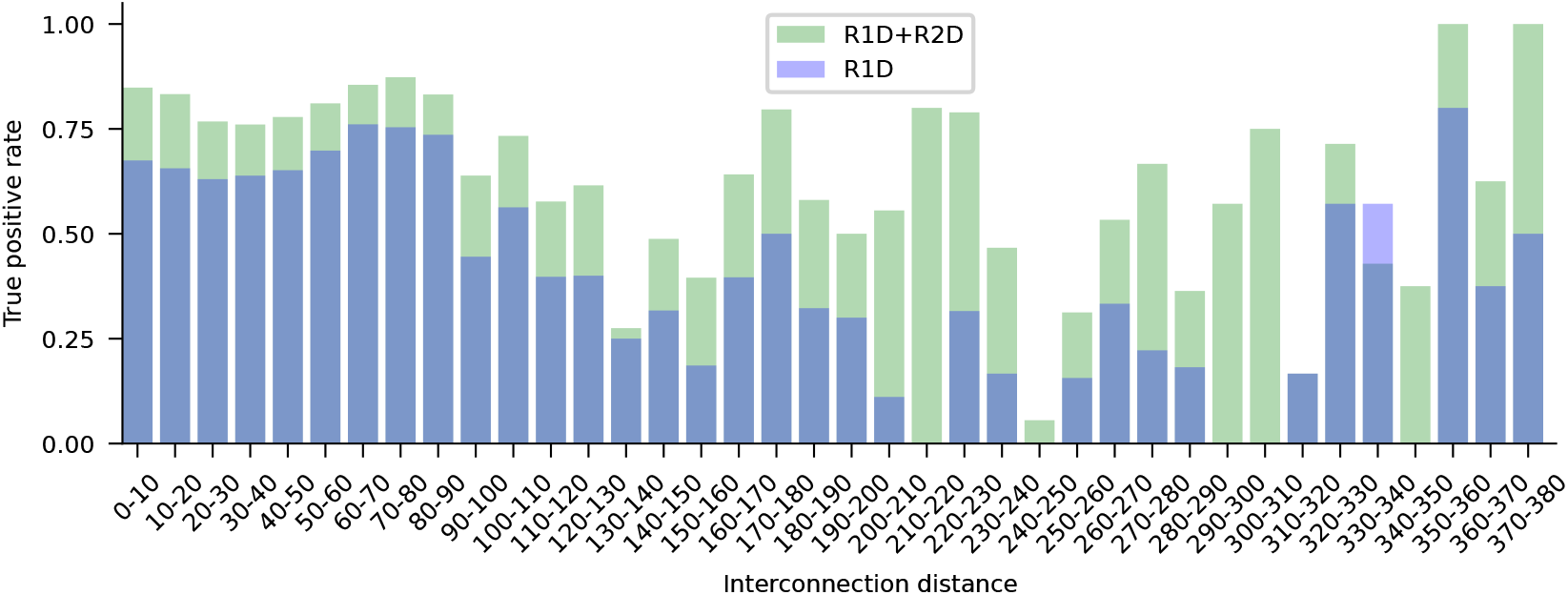
True positive rate for each interaction distance, comparing the model with only the first stage and both stages. Resnet based model with only the first stage (R1D) and both stages (R1D+R2D).

This shows that in fact the sincFold 2D stage improves the learning of long range dependencies. Moreover, a very interesting property of ResNet blocks is that when there are many blocks available, the model is capable of automatically selecting how many of them are really necessary, skipping the non-necessary blocks during training. This reduces the learnable parameters while helping to learn more relevant features.

Using the best performing sincFold architecture (R1D+R2D), we performed a hyperparameter space search in the Ablation dataset, exploring: batch size, learning rate, the use of learning rate schedule, weights for the loss components *λ*_*β*_, *λ*_1_, architecture of the 1D-ResNet stage (kernel size and dilation, number of filters and number of layers) and the 2D-ResNet stage (kernel size, number of filters, bottleneck size and number of layers). Parameters were explored randomly [6], and the best configuration was selected for the next experiments (Supplementary Material Figure S1).

### 4.2 Performance according to test-train structural distance on random partitions

Figure 4 shows the comparative results among classical folding methods (RNAfold, RNAstructure, ProbKnot, IPKnot, LinearPartition-V, LinearFold-V, LinearPartition-C and LinearFold-C), the hybrid method MXfold2, DL based methods (UFold and REDfold) and the proposed sincFold in terms of *F*_1_ for 5-fold cross-validation on the RNAstralign dataset. All DL methods were trained and evaluated from scratch with the same dataset partitions on cross-validation. It can be seen that all classical methods have a performance between 0.633 and 0.712 of *F*_1_. MXFold2 combines DL and thermodynamic models and achieves better performance (median *F*_1_ = 0.907). DL methods show even better scores, UFold reaches a median *F*_1_ = 0.966 and REDfold arrives at median *F*_1_ = 0.976.

**Figure 4:**
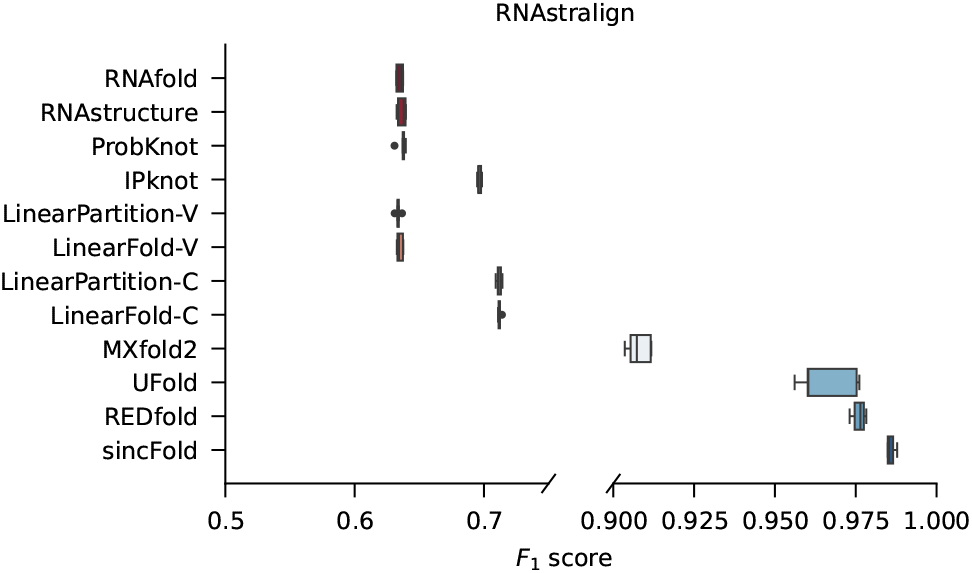
Comparative results among classical folding methods, DL-based folding methods and sincFold, for a 5-fold cross-validation on the RNAstralign dataset. Horizontal scale was adjusted to improve visualization.

The proposed method, sincFold, achieves *F*_1_ = 0.986. The variance of our method is very small, and the box is not overlapped with the performance of the other DL methods.

Figure 5 shows the comparative results among classical folding methods, hybrid method, DL methods and the proposed sincFold, in terms of *F*_1_ for the ArchiveII dataset. As in the previous result, classical methods have a median performance below *F*_1_ = 0.620. In this case, MxFold2 achieves *F*_1_ = 0.738, UFold has a median *F*_1_ = 0.855 and REDfold arrives at *F*_1_ = 0.831. The proposed method sincFold achieves the highest median *F*_1_ = 0.913. It can be seen that in both datasets, our proposal achieves a significantly better performance than classical methods and state-of-the-art DL methods.

**Figure 5:**
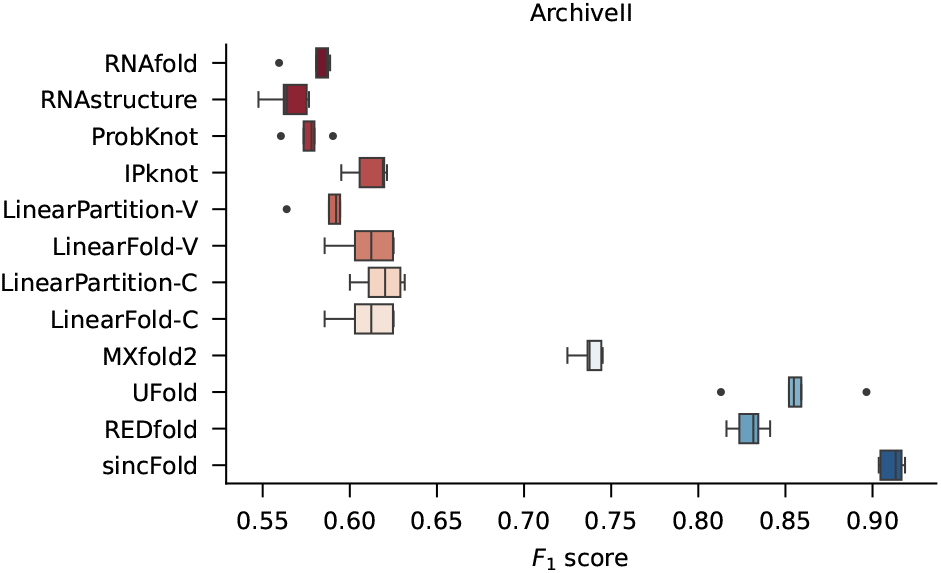
Comparative results among classical folding methods, DL-based folding methods and sincFold, for cross-validation on the ArchiveII dataset.

The detailed performance for each method according to the lengths of the test sequences, from shorter (left) to longer sequences (right) are analyzed in Figure 6. Light blue bars indicate the proportion of each bin of lengths in the dataset. Here it can be seen that for shorter sequences, all methods have average performance above *F*_1_ = 0.60, being particularly good at this task all DL methods, with *F*_1_ *>* 0.90. As sequence length is increased, classical methods lower performance while DL methods are less affected, maintaining *F*_1_ *>* 0.75 in most cases and being sincFold always the best for long sequences, achieving *F*_1_ = 0.85 for sequences between 300 and 400 nucleotides. At the extreme, for sequences longer than 400 nucleotides, sincFold is still better than other methods despite the few examples available to learn from, achieving a median *F*_1_ = 0.74, which is superior to the average performance of classical methods in the shortest sequences.

**Figure 6:**
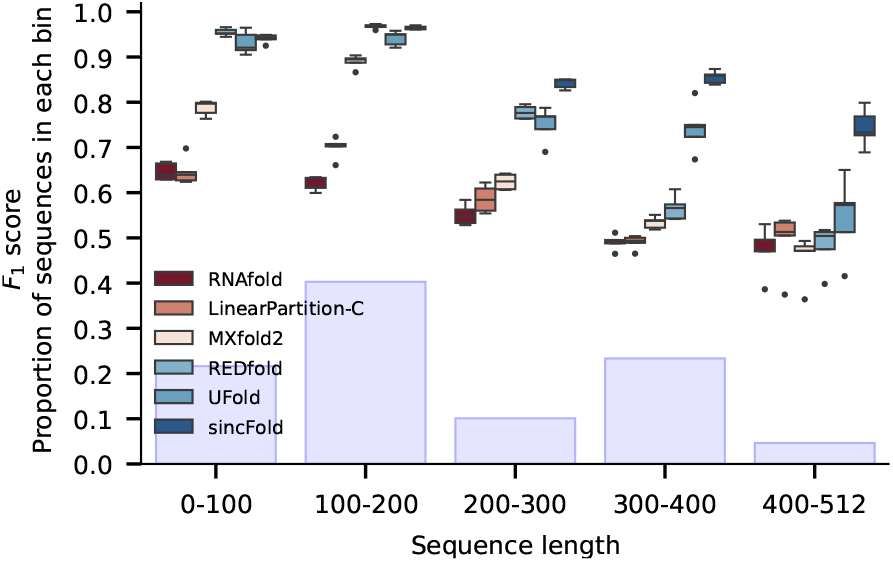
Detailed *F*_1_ performance for each method according to the mean lengths of the sequences, from shorter (left) to longer (right) sequences.

It is well-known that at random partitions there can be sequences with high similarity between training and testing partitions, thus methods can show overly optimistic results. To have more insights on the sincFold performance in this regard, we report comparative results by analyzing the sequences following the secondary structure distance between test and train partitions, as shown in Figure 7. Instead of just making one single partition with a certain sequence identity level, we have analyzed a full range of structural similarities. The distance between two structures was computed using RNAdistance from the ViennaRNA package [25] as explained in Section 3.3. For each test sequence, the test-train distance was defined as the minimum structural distance between this test sequence and all the sequences in its corresponding training fold. Then, test sequences with similar structural distance were grouped into bins to obtain the x-axis in the figure. Ranges of structural distances are presented from large (very-hard) test-train distances (left) down to low (very-easy) test-train distances (right). Light blue bars indicate the proportion of test sequences in each bin of structural distances.

**Figure 7:**
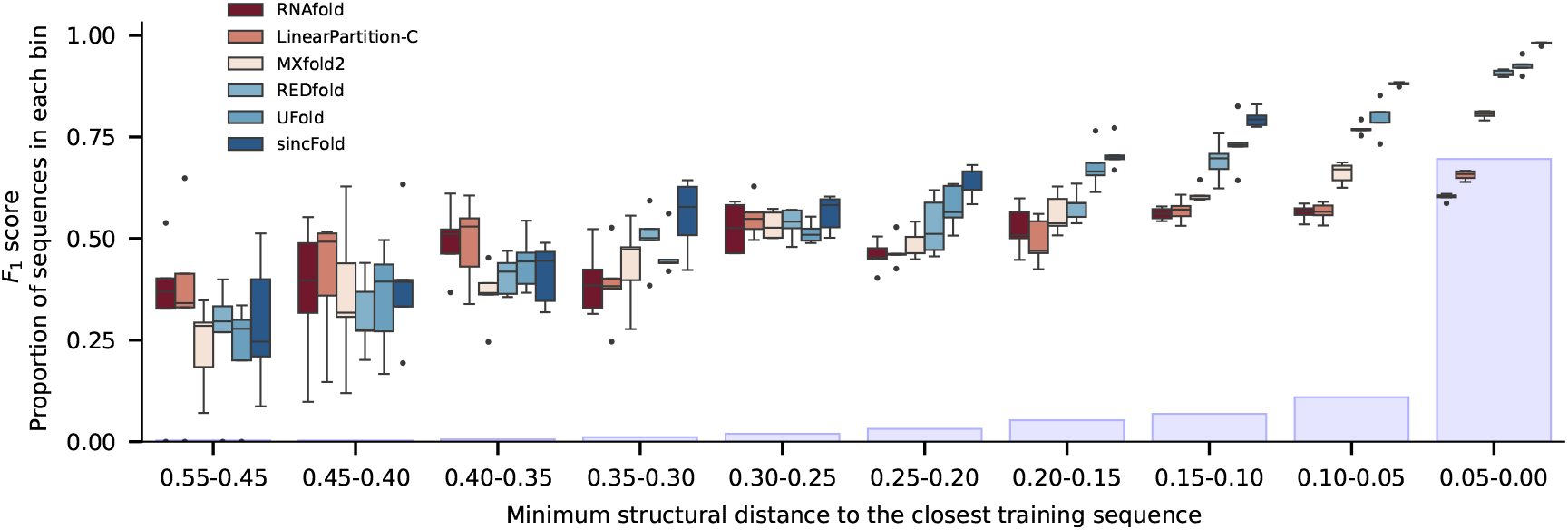
Mean *F*_1_ scores for each method according to test-train structural distance, from large distances (left) to low distances (right). Bars indicate the proportion of each bin in the dataset.

It can be clearly seen that as the test-train distance diminishes, all methods (including the classical, non-learnable ones) improve performance. This also makes evident that structures on the left side are really harder to predict, even for classical models that are not trained and thus they are agnostic to the test-train structural distances. For the 23 structures in the first 2 bins of distances all methods have median *F*_1_ *<* 0.50. It can be seen that for distances between 0.40 and 0.25, both classical and DL methods have again a low performance *F*_1_ *∈* (0.30, 0.60). In the middle cases, from 0.25 to 0.20 distance, DL methods are slightly better than classical ones. Finally, for the lowest test-train structural distances (*<* 0.15), DL methods are clearly better for RNA secondary structure prediction, being sincFold the best method in all cases, improving classical methods from a distance of 0.25 and all other trainable methods from 0.20. These trends can be explained by two facts. First, the abundance of structure samples benefits DL models more than the classical ones because the former have more cases to learn from. Besides, the benefits for the classical methods are indirect since they do not learn, but were developed looking at the most abundant or popular structures that therefore better fit the thermodynamic models. Secondly, based on the advantage of structure abundance for data-driven approaches, sincFold is the one that best takes advantage of the ability to learn from more distant samples, regardless of how much is known about the thermodynamics of the molecules. This is evident even when distance is around 0.25 and thus far from overfitting from training samples.

### 4.3 Homology-aware validation

For a deeper performance analysis of the methods considering homology between training and testing partitions, in this section we performed experiments with a more rigorous control of homology, instead of using random partitions. Table 1 shows the results of testing models in a non-redundant set of RNA sequences at 80% sequence-identity cutoff (TR0-TS0 partitions). The table reports comparative results on the TS0 test set according to *F*_1_, *WL* and *INF* metrics. In all cases, the three metrics are consistent and show that sincFold is the best method to predict RNA structures with low homology to the training set, and there is almost a 10% performance gap with the classical methods.

**Table 1:** Performance for methods on bpRNA. The TR0 partition is used for training and the TS0 partition for testing.

| Method | $F_1$ | $WL$ | $INF$ |
| --- | --- | --- | --- |
| RNAfold | 0.508 | 0.681 | 0.520 |
| LinearPartition-C | 0.547 | 0.737 | 0.556 |
| MXfold2 | 0.502 | 0.736 | 0.509 |
| REDfold | 0.621 | 0.799 | 0.626 |
| UFold | 0.580 | 0.743 | 0.590 |
| sincFold | 0.631 | 0.827 | 0.641 |

For benchmarking inter-family performance, a family-fold cross-validation in the ArchiveII dataset was performed. That is, one family is left out for testing per cross-validation fold and the rest of the families are used for training as in [45]. This eliminates most of the homology to the training set, providing a hard measure of performance and, thus, allows estimating future performance on novel RNAs that do not belong to any well-known family.

Table 2 shows the average inter-family performance comparison with several metrics between sincFold and other classical, hybrid and pure DL methods for RNA secondary structure prediction. It can be seen that all the three metrics used, *F*_1_ score, *WL* and *INF* metric, consistently indicate that sincFold is the best DL model to predict novel RNA structures of families of RNA never seen during training. As seen previously, performance of the hybrid method is in-between classical and DL methods. Obviously classical methods obtain the best performance here since they do not fully comply with the cross-family validation. This is because they use constraints and thermodynamic parameters that have been experimentally determined from the hairpin loops and other important structures that were most frequently found in most of the RNA families in this dataset [2, 3, 18, 28, 33]. In terms of *F*_1_, the difference between the best DL method (sincFold) and the best classical method (LinearPartition-C) is 0.191, and 0.193 in the case of *INF* . However, looking at the WL metric this gap is much lower, being 0.098. This suggests that when measuring the graph structural information of the predictions, DL and classical methods are close in performance for inter-family validation.

**Table 2:**
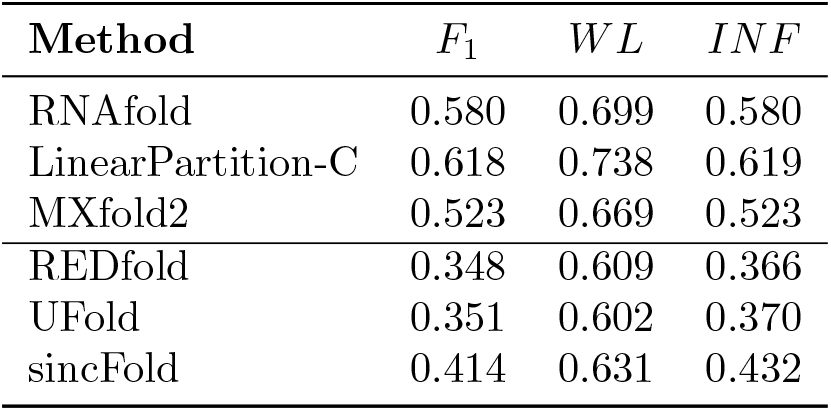
Inter-family performance comparison in ArchiveII, between sincFold and classical, hybrid and DL methods.

| Method | $F_1$ | $WL$ | $INF$ |
| --- | --- | --- | --- |
| RNAfold | 0.580 | 0.699 | 0.580 |
| LinearPartition-C | 0.618 | 0.738 | 0.619 |
| MXfold2 | 0.523 | 0.669 | 0.523 |
| REDfold | 0.348 | 0.609 | 0.366 |
| UFold | 0.351 | 0.602 | 0.370 |
| sincFold | 0.414 | 0.631 | 0.432 |

Table 3 shows the detailed performance of DL methods for each family in the ArchiveII dataset. Full results for all methods and all measurements for each family can be found in the Supplementary Material, Table S1.

**Table 3:**
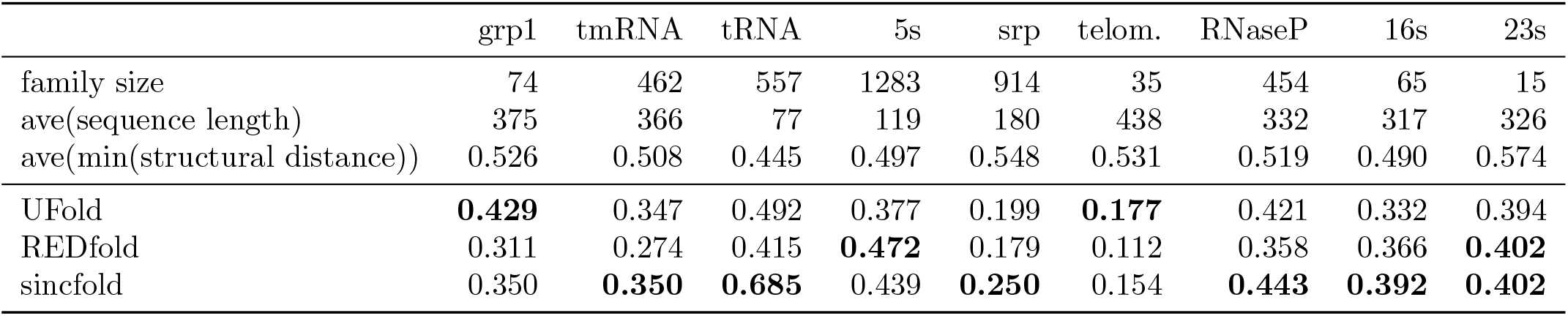
Inter-family performance detail of the *F*_1_ score in ArchiveII for each RNA family in the comparison between sincFold and other DL RNA secondary structure prediction methods.

The 9 RNA families are characterized in terms of number of samples, average length, structural distance and sequential distance to the other families. It can be seen that when the grp1 family is used as the test set, all methods have a moderate to low performance. The best DL method here achieves *F*_1_ = 0.429. This is a family with a very low number of examples, which have a mean sequence length that is longer than most of the other families, with a moderate structural distance to the other families and high sequence distance to the rest of the dataset. In the case of the tmRNA family, all methods have low performance as well, but here both sincFold and UFold achieve the best result. In spite of having more testing examples (and thus the training set is much smaller), the characteristics of this family are similar to grp1 regarding structure and sequence distance to the training set, while sincFold achieves a similar results. The tRNA family is the one with the lowest sequence length, having a large number of examples. In this case, while the other DL methods have low performance, here sincFold achieves a performance of *F*_1_ = 0.685 that is very close to the performance of many classical methods (Table S1). The srp and telomerase testing families are the hardest ones. The srp family, which is indeed very different from the rest of the families regarding sequence distance and structural distance, is better predicted by sincFold. The telomerase family has a very low number of samples, which are indeed very different from the rest of the families regarding mean sequence length (those are the longest sequences, almost double the average). In the case of the 16s and 23s families, also sincFold provides the best predictions.

In summary, sincFold shows improved performance in comparison with the other DL methods in the prediction of tmRNA, tRNA, srp, RNAseP, and 16s families; that is 5 out of 9 families. This is further evidence of the improved generalization capability that sincFold provides relative to the state-of-the-art DL methods.

## 5. Conclusions

In this work, we presented sincFold, an end-to-end deep learning model that can accurately predict the secondary structure from a RNA sequence without requiring multi-sequence alignments, or any other pre-processing of the input sequences. Local and distant relationships can be learnt effectively using a sequential 1D-2D architecture. Based on ResNet blocks, bottlenecks layers and a 1D-to-2D projection, it has proven to be better suited to identify structures that might defy traditional modeling, while reducing the effective number of trainable parameters. We show that sincFold outperforms other methods even with moderate structural distances between train and testing sequences. Results also show that sincFold, thanks to its capability for capturing a wide range distances in interactions, is significantly better than all other methods for the secondary structure prediction also in longer ncRNA sequences (more than 200 nucleotides). In an inter-family evaluation, sincFold performed better than other state-of-the-art DL approaches, showing that RNA structure predictions can still be improved with trainable methods.

## Competing interests

No competing interest is declared.

## Funding

We would like to thank to the AWS Cloud Credit for Research, the Ministerio de Producción, Ciencia y Tecnología, Santa Fe (PEICID-2022-075) and the Agencia Nacional de Promoción de la Investigación, el Desarrollo Tecnológico y la Innovación (Ciencia y tecnología contra el hambre) for their support for this work. This work was supported by ANPCyT (PICT 2018 3384, PICT 2022 0086) and UNL (CAI+D 2020 115). We also thank Nvidia for donating Titan XP GPUs used in the research.

## Footnotes

1 Source code available at https://github.com/sinc-lab/sincFold

2 Web-demo available at https://sinc.unl.edu.ar/web-demo/sincFold

## Notes

https://github.com/sinc-lab/sincFold

### Competing Interest Statement

The authors have declared no competing interest.

### Summary of Updates

Text in general was improved, adding details. Datasets and results were added to improve model validation, specially section 4.3

