## Supplementary material for "sincFold: end-to-end learning of short- and long-range interactions in RNA secondary structure"

### 1 Hyperparameter search

The exploration of hyperparameters on the Ablation dataset resulted in a set of 4021 runs with different parameters and an average $F_1$ score. A random forest regressor was trained on this data to explore the hyperparameter effect (predictability) on $F_1$. Using the mean decrease of impurity [1] (Figure S1) the hyperparameters batch size, learning rate (LR), use of learning rate schedule, weights $\lambda_\beta$, $\lambda_{L_1}$ and the architecture of the ResNet stages were analyzed and selected iteratively. The LR, ResNet2D kernel size, bottleneck filters on the first ResNet layer (Bottleneck$_1$), batch size and number of layers in the ResNet2D resulted in the most critical parameters, in the sense that poor choices may result in strong loss of performance (Figure S2). Optimal LR found was 0.001 and batch size less than 8. In the ResNet2D stage, 2 layers with kernel size 7 were the best choice. Finally, it was better to use a small kernel of size 3 in ResNet1D.

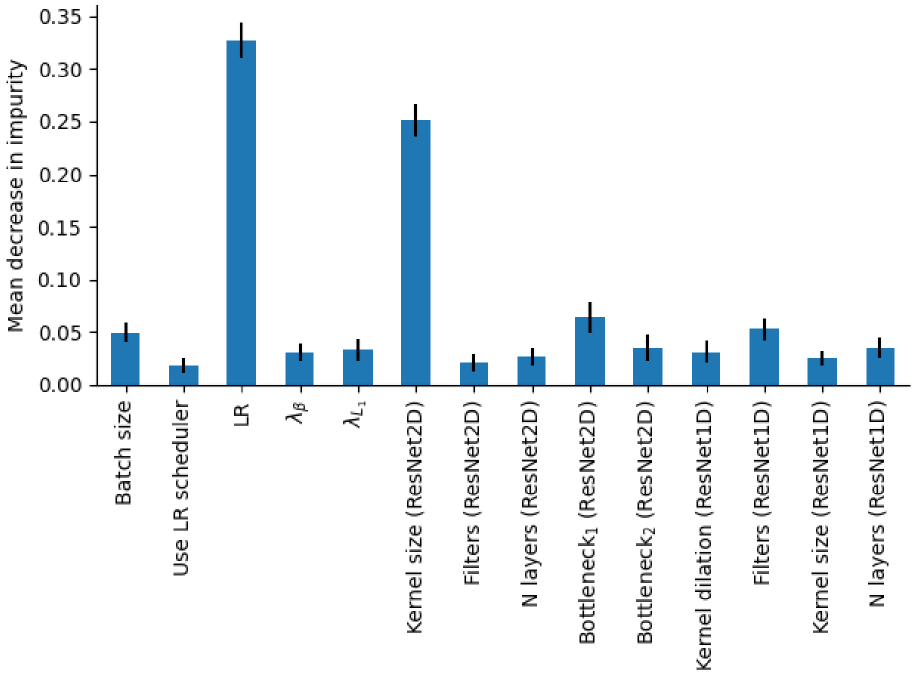


Figure 1: Hyperparameter importance based on the mean decrease in impurity of the random forest regressor. Higher values indicate more predictability (correlation) between hyperparameter change and resulting $F_1$ score.

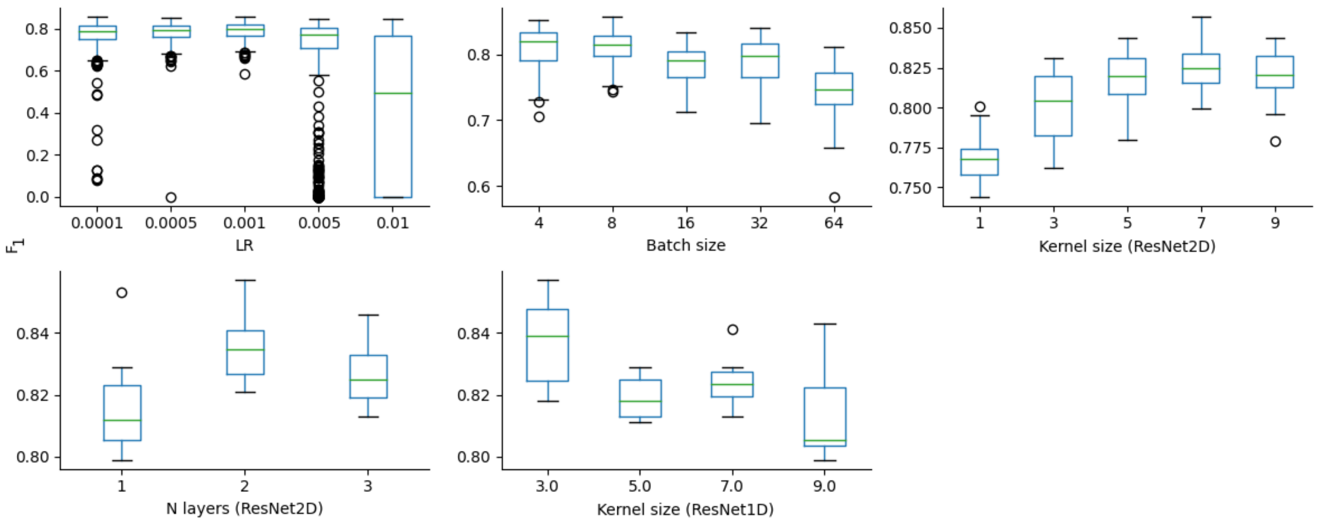


Figure 2: Impact on $F_1$ score of the critical hyperparameters of sincFold model.

### 2 Inter-family results

Table 1: All metrics for the inter-family experiment.

| | | grp1 | tmRNA | tRNA | 5s | srp | telom. | RNaseP | 16s | 23s |
|---|---|---|---|---|---|---|---|---|---|---|
| family size | | 74 | 462 | 557 | 1283 | 914 | 35 | 454 | 65 | 15 |
| ave($L$) | | 375 | 366 | 77 | 119 | 180 | 438 | 332 | 317 | 326 |
| ave(min(structural distance)) | | 0.526 | 0.508 | 0.445 | 0.497 | 0.548 | 0.531 | 0.519 | 0.490 | 0.574 |
| $F_1$ | | | | | | | | | | |
| Classical methods | RNAfold | 0.540 | 0.419 | 0.677 | 0.614 | 0.593 | 0.465 | 0.527 | 0.532 | 0.694 |
| | RNAstructure | 0.494 | 0.409 | 0.690 | 0.584 | 0.568 | 0.443 | 0.532 | 0.533 | 0.663 |
| | IPknot | 0.587 | 0.394 | 0.705 | 0.692 | 0.588 | 0.517 | 0.556 | 0.580 | 0.689 |
| | LinearPartition-C | 0.592 | 0.370 | 0.699 | 0.718 | 0.589 | 0.516 | 0.560 | 0.594 | 0.718 |
| | LinearPartition-V | 0.545 | 0.404 | 0.658 | 0.632 | 0.596 | 0.453 | 0.554 | 0.550 | 0.717 |
| | LinearFold-C | 0.545 | 0.390 | 0.673 | 0.726 | 0.589 | 0.490 | 0.497 | 0.565 | 0.657 |
| | LinearFold-V | 0.545 | 0.390 | 0.673 | 0.726 | 0.589 | 0.490 | 0.497 | 0.565 | 0.657 |
| | ProbKnot | 0.535 | 0.415 | 0.680 | 0.599 | 0.573 | 0.465 | 0.572 | 0.558 | 0.692 |
| Hybrid | MXfold2 | 0.459 | 0.411 | 0.531 | 0.555 | 0.561 | 0.363 | 0.482 | 0.520 | 0.594 |
| DL | UFold | 0.429 | 0.347 | 0.492 | 0.377 | 0.199 | 0.177 | 0.421 | 0.332 | 0.394 |
| | REDfold | 0.311 | 0.274 | 0.415 | 0.472 | 0.179 | 0.112 | 0.358 | 0.366 | 0.402 |
| | sincfold | 0.350 | 0.350 | 0.685 | 0.439 | 0.250 | 0.154 | 0.443 | 0.392 | 0.402 |
| $INF$ | | | | | | | | | | |
| Classical methods | RNAfold | 0.541 | 0.420 | 0.677 | 0.613 | 0.593 | 0.471 | 0.529 | 0.534 | 0.694 |
| | RNAstructure | 0.496 | 0.410 | 0.689 | 0.584 | 0.568 | 0.449 | 0.533 | 0.535 | 0.664 |
| | IPknot | 0.588 | 0.396 | 0.707 | 0.692 | 0.589 | 0.520 | 0.560 | 0.582 | 0.691 |
| | LinearPartition-C | 0.592 | 0.373 | 0.700 | 0.718 | 0.590 | 0.520 | 0.563 | 0.595 | 0.719 |
| | LinearPartition-V | 0.547 | 0.406 | 0.658 | 0.632 | 0.597 | 0.459 | 0.556 | 0.553 | 0.718 |
| | LinearFold-C | 0.546 | 0.393 | 0.678 | 0.728 | 0.590 | 0.494 | 0.500 | 0.567 | 0.659 |
| | LinearFold-V | 0.546 | 0.393 | 0.678 | 0.728 | 0.590 | 0.494 | 0.500 | 0.567 | 0.659 |
| | ProbKnot | 0.536 | 0.416 | 0.681 | 0.598 | 0.573 | 0.470 | 0.573 | 0.560 | 0.692 |
| Hybrid | MXfold2 | 0.459 | 0.412 | 0.530 | 0.554 | 0.561 | 0.365 | 0.483 | 0.521 | 0.593 |
| DL | UFold | 0.434 | 0.350 | 0.515 | 0.408 | 0.209 | 0.177 | 0.438 | 0.345 | 0.415 |
| | REDfold | 0.317 | 0.278 | 0.442 | 0.498 | 0.189 | 0.114 | 0.374 | 0.377 | 0.421 |
| | sincfold | 0.362 | 0.356 | 0.696 | 0.475 | 0.260 | 0.160 | 0.449 | 0.397 | 0.430 |
| $WL$ | | | | | | | | | | |
| Classical methods | RNAfold | 0.673 | 0.591 | 0.765 | 0.717 | 0.712 | 0.610 | 0.666 | 0.685 | 0.773 |
| | RNAstructure | 0.642 | 0.582 | 0.776 | 0.697 | 0.695 | 0.598 | 0.669 | 0.684 | 0.758 |
| | IPknot | 0.716 | 0.599 | 0.781 | 0.771 | 0.712 | 0.658 | 0.695 | 0.733 | 0.771 |
| | LinearPartition-C | 0.725 | 0.592 | 0.791 | 0.796 | 0.719 | 0.659 | 0.701 | 0.750 | 0.793 |
| | LinearPartition-V | 0.672 | 0.577 | 0.750 | 0.725 | 0.711 | 0.596 | 0.681 | 0.692 | 0.786 |
| | LinearFold-C | 0.707 | 0.606 | 0.783 | 0.809 | 0.724 | 0.644 | 0.668 | 0.739 | 0.763 |
| | LinearFold-V | 0.707 | 0.606 | 0.783 | 0.809 | 0.724 | 0.644 | 0.668 | 0.739 | 0.763 |
| | ProbKnot | 0.670 | 0.586 | 0.760 | 0.709 | 0.699 | 0.613 | 0.691 | 0.696 | 0.766 |
| Hybrid | MXfold2 | 0.632 | 0.599 | 0.673 | 0.690 | 0.692 | 0.562 | 0.644 | 0.688 | 0.720 |
| DL | UFold | 0.644 | 0.579 | 0.675 | 0.628 | 0.517 | 0.528 | 0.636 | 0.616 | 0.623 |
| | REDfold | 0.595 | 0.566 | 0.658 | 0.669 | 0.518 | 0.541 | 0.612 | 0.646 | 0.628 |
| | sincfold | 0.610 | 0.582 | 0.783 | 0.653 | 0.538 | 0.579 | 0.631 | 0.627 | 0.631 |
